# Targeting a Granulocytic–Microbiome Axis Reverses Glioblastoma Progression via Intranasal Cannabidiol

**DOI:** 10.64898/2026.07.12.737962

**Authors:** Lei P Wang, Bidhan Bhandari, Sahar Emami Naeini, James T Earwood, Brendan Marshall, Chandramohan Wakade, Jack C Yu, Ali S Arbab, Évila Lopes Salles, Babak Baban

## Abstract

Mucosal cannabidiol formulations are known regulators of the glioblastoma microenvironment, yet the underlying origin point triggering this stroma-remodeling efficacy remains entirely unknown. Here, by mapping innate cell trafficking pathways, we define a novel baseline neuro-immune-microbiome axis in orthotopic glioblastoma, characterized by diverse microbial communities, likely seeded via blood-brain barrier disruption, paired with dense infiltration of host mast cells and mature, crystalloid-containing eosinophils. Localized intranasal administration of a synthetic cannabidiol formulation achieved striking therapeutic efficacy, driving dramatic tumor regression. Mechanistically, high-throughput 16S rRNA sequencing and quantitative flow cytometry revealed this progression was subverted by taming the tumor ecosystem; cannabidiol restricted chaotic microbial diversity, selectively filtering the landscape toward Delftia and depleting Archaea, while simultaneously suppressing hyper-inflammatory host mast cell and eosinophil populations. This study builds upon established innate trafficking frameworks to present the first therapeutically targetable stromal-microbial axis in neuro-oncology.

## Introduction

Glioblastoma (GBM) remains among the most lethal malignancies of the central nervous system, characterized by a highly infiltrative growth pattern and a profoundly volatile, immunosuppressive tumor microenvironment (TME). Despite aggressive surgical and chemoradiotherapeutic interventions, therapeutic evasion remains the clinical norm ^1-3^. This resistance is driven by the TME’s ability to sequester the tumor from systemic surveillance while fostering localized, hyper-inflammatory niche factors that drive progression. While the central nervous system was historically viewed as a sterile sanctuary, the “sterile brain” paradigm has been recently dismantled by landmark characterizations identifying distinct intratumoral microbial signatures within high-grade gliomas ^4^. These microbial communities are likely seeded via blood-brain barrier (BBB) disruption, establishing a complex, uncharacterized stromal-microbial ecosystem. However, the precise baseline interaction between these resident intratumoral microbes and host innate immune cells, and whether this axis can be pharmacologically manipulated, remains entirely unknown.

An emerging yet comparatively underrecognized component of the glioblastoma immune landscape is the granulocytic compartment, particularly infiltrating host mast cells (MCs) and eosinophils. Although these cells have been studied extensively in allergic disorders and peripheral inflammation, their roles within glioblastoma remain incompletely understood ^5-9^. Accumulating evidence suggests that mast cells and eosinophils infiltrate glioblastoma in a grade-dependent manner, preferentially localizing within perivascular and stromal niches where they regulate extracellular matrix remodeling, vascular integrity, cytokine signaling, and the maintenance of perivascular glioma stem cell niches ^9-11^. Rather than functioning solely as classical inflammatory effectors, these granulocyte populations may actively remodel the glioblastoma immune ecosystem by promoting immune dysregulation, tissue plasticity, and tumor progression^9,12-14^ .

Our previous studies established that localized mucosal cannabidiol delivery suppresses glioblastoma progression through comprehensive remodeling of the tumor microenvironment, including modulation of inflammatory signaling, innate immune composition, and tumor-supportive cellular interactions^15,16^. We subsequently demonstrated that early immunological conditioning of the glioblastoma microenvironment can influence disease trajectory and provided proof-of-concept evidence that peripheral innate lymphoid cells actively home to the glioblastoma niche, underscoring the dynamic communication between the peripheral immune system and the central nervous system tumor ecosystem ^17,18^. Building upon these findings, we hypothesized that this immune remodeling extends beyond host immunity alone to encompass the broader biological ecosystem of glioblastoma, including its resident microbial landscape and infiltrating granulocyte populations.

In the present study, we define a previously unrecognized neuro-immune-microbial axis within orthotopic glioblastoma. Using high-resolution transmission electron microscopy, immunofluorescence, quantitative flow cytometry, and 16S rRNA sequencing, we identify a complex baseline ecosystem characterized by coexisting microbial communities together with prominent infiltration of mature eosinophils and activated mast cells. We further demonstrate that non-invasive intranasal administration of a synthetic cannabidiol (CBD) isolate profoundly remodels this ecosystem by simultaneously reshaping the tumor-associated microbiome and attenuating host granulocytic inflammation, resulting in marked suppression of glioblastoma progression. Collectively, these findings identify an unrecognized interface between the tumor microbiome and innate immunity and provide a mechanistic framework through which targeted cannabinoid therapy may restore immune-microbial homeostasis within the glioblastoma microenvironment.

## Materials and Methods

### Orthotopic Glioblastoma Mouse Model

All animal procedures and surgical workflows were formally reviewed and approved by the Institutional Animal Care and Use Committee (IACUC) at Augusta University. For intracranial tumor modeling, a total of 18 male and female C57BL/6 mice (n = 9 animals per experimental arm, pooled across three distinct, independent cohorts) were utilized. Intracranial niches were established using firefly luciferase-expressing GL261 cells (GL261-Luc2) obtained from the Georgia Cancer Center Cell Repository, adopting the immunocompetent system previously validated and published by our laboratory ^18^. Surgical anesthesia was induced via 3% isoflurane inhalation and maintained continuously between 1.5% and 2% throughout the stereotaxic procedure. Animals received a localized microinjection of 3 × 10^4^ GL261-Luc2 cells suspended in 3 µL of sterile phosphate-buffered saline, targeted precisely to the right striatal coordinates to model a syngeneic, limited-infiltration malignant environment. Longitudinal engraftment and subsequent baseline kinetic growth were monitored non-invasively via optical bioluminescence imaging (IVIS Spectrum). At the definitive experimental endpoint, cohorts were humanely sacrificed via carbon dioxide inhalation followed by secondary cervical dislocation in strict accordance with AVMA protocols.

### Intranasal Cannabidiol Formulation and Administration

Therapeutic interventions utilized an analytical-grade, synthetic, Δ^9^-THC-free cannabidiol (CBD) isolate manufactured by Purysis and procured commercially via Restek (Bellefonte, PA, USA). The crystalline isolate was dissolved and homogenized in a biocompatible, mucosal-adhesive polymeric vehicle optimized specifically for targeted delivery to the central compartment via the nasal mucosa. For the intervention cohort, mice received daily localized intranasal administrations consisting of 25 µL of the CBD formulation (45 mg/mL) per nostril, yielding a fixed dose of 2.25 mg CBD per mouse per day. To promote mucosal retention and minimize the risk of aspiration, the formulation was instilled slowly in sequential micro-droplets to conscious animals. Vehicle-control cohorts received an identical volume of the mucoadhesive polymeric vehicle alone. Treatments commenced on post-implantation day 9 following tumor establishment and were maintained daily for 12 consecutive days until the 3-week experimental endpoint.

### 16S rRNA Microbial Sequencing and Bioinformatics

To characterize the localized intratumoral microbiome, intracranial glioblastoma tissue cores were surgically micro-dissected at the experimental endpoint. To ensure complete elimination of environmental or procedural microflora contamination, all tissue harvesting was executed under strict aseptic conditions using sterile, DNA-free instruments. Collected tumor samples were immediately flash-frozen and preserved at −80°C until processing. Total genomic DNA isolation from the glioblastoma cores was executed in-house utilizing a specialized genomic DNA extraction kit (QIAGEN, Germantown, MD, USA) according to the manufacturer’s protocol, leveraging mechanical homogenization optimized for low-biomass tissue matrices and tough microbial cell walls.

Purified genomic DNA extracts were transferred to Novogene Corporation (Durham, NC, USA) for amplicon library construction and high-throughput sequencing. The bacterial 16S rRNA gene V3–V4 hypervariable regions were targeted and amplified via polymerase chain reaction (PCR) utilizing barcoded locus-specific primers prior to multiplexed sequencing on an Illumina platform. Raw sequencing outputs were resolved through the QIIME2 bioinformatic pipeline. Alpha-diversity (Shannon entropy index) and beta-diversity metrics were calculated alongside reference-based taxonomic classification against the SILVA reference database. Characterized microbial community signatures, including specific intratumoral taxonomic filtering and Archaea depletion, were statistically evaluated and visualized across experimental cohorts.

### Flow Cytometric Analysis of Infiltrating Granulocytes

Quantification of intratumoral immune cell populations was executed via high-dimensional flow cytometry following single-cell isolation protocols previously established and validated by our laboratory ^17^. Briefly, harvested glioblastoma tissue cores underwent mechanical dissociation followed by enzymatic digestion in a collagenase/DNase cocktail at 37°C to yield uniform cellular suspensions. Prior to surface marker labeling, nonspecific binding was restricted utilizing an anti-CD16/CD32 Fc-receptor blocking antibody. To precisely characterize the infiltrating host granulocyte architecture for the first time in this model, cells were incubated with a novel, fluorophore-conjugated antibody panel optimized specifically for myeloid lineages. Infiltrating host mast cells were rigorously defined and gated using the multi-marker strategy CD45^+^FcεRI^+^CD117^+^. Concurrently, host eosinophil populations were resolved and quantified as CD45^+^CD11b^+^Siglec-F^+^SSC^high^. High-dimensional acquisition was performed on a calibrated flow cytometer, and downstream cluster analysis and absolute cell profiling were conducted using FlowJo software.

### Fluorescence-Based Immunohistochemical Analysis and Spatial Mapping

For structural localization and spatial mapping within the intracranial stroma, immunofluorescence staining was carried out on paraffin-embedded brain tumor tissue sections adopting protocols previously established and validated by our laboratory ^17^. Following deparaffinization and antigen retrieval boundaries, tissue sections were incubated overnight with a cocktail of fluorophore-conjugated primary antibodies (BioLegend, San Diego, CA, USA, unless stated otherwise) optimized for multi-marker granulocytic identification. Infiltrating host mast cells were phenotypically mapped using anti-CD117 and anti-Tryptase antibodies, while host eosinophils were structurally defined using anti-CD45, anti-Siglec-F, and anti-CD193 antibodies. Nuclear counterstaining was executed utilizing DAPI (4′,6-diamidino-2-phenylindole) to facilitate deep cellular visualization within the perivascular and core tumor niches. High-resolution spatial profiles and multi-channel fluorescent images were captured using a Zeiss fluorescence microscope ecosystem (Zeiss USA, White Plains, NY, USA). Quantitative digital image analysis was subsequently conducted by defining standardized regions of interest (ROIs) with the selection mechanics in Adobe Photoshop CS4 Extended (version 11.0; Adobe Systems, San Jose, CA, USA). Within these designated spatial boundaries, the integrated density (sum of all pixel intensities) and mean gray value (quantifying relative fluorescence brightness) were recorded systematically to evaluate comparative marker expression and architectural shifts across experimental cohorts.

### Transmission Electron Microscopy (TEM)

To achieve nanometer-scale ultrastructural resolution of host granulocyte architecture and specific activation states, fresh, millimeter-sized glioblastoma tissue blocks were processed following electron microscopy protocols previously established and validated by our laboratory ^19,20^. Tissues were immediately immersed in ice-cold primary fixative comprising 2% glutaraldehyde and 2% paraformaldehyde buffered in 0.1 M sodium cacodylate (pH 7.4). Following baseline fixation, specimens were post-fixed in 1% osmium tetroxide, systematically dehydrated through a graded ethanol series, and embedded in Eponate 12 resin blocks.

Ultra-thin sections (70–80 nm) were cut using an ultramicrotome, mounted onto copper grids, and contrast-stained with uranyl acetate and lead citrate. Ultrastructural imaging was executed on a transmission electron microscope operating at an accelerating voltage of 80 kV. This high-resolution approach was leveraged to visually document and differentiate intact versus degranulating host mast cell profiles alongside mature host eosinophils packed with distinct crystalloid-containing secondary granules.

### Statistical Analysis

Quantitative datasets are expressed as mean ± standard error of the mean (SEM). Group sample sizes (n = 9 mice per arm, generated from three independent experimental replicates) were prospectively calculated via power analysis to ensure 80% statistical power (α = 0.05) for detecting a minimum 30% variance in overall tumor burden. Longitudinal bioluminescence measurements reflecting intracranial tumor progression were statistically evaluated using a two-way analysis of variance (ANOVA) followed by Tukey’s multiple comparisons post hoc test. Animal survival curves were plotted via Kaplan–Meier analysis, and statistical disparities between cohorts were determined using the log-rank (Mantel–Cox) test. Thresholds for statistical significance were stringently defined as *P < 0.05, * *P < 0.01, and * * *P < 0.001. All computational modeling and statistical evaluations were conducted utilizing GraphPad Prism version 9.0 (GraphPad Software, Boston, MA, USA).

## Results

### Intranasal CBD Restricts Glioblastoma Expansion In Vivo

To evaluate our intranasal CBD formulation against aggressive intracranial tumors, we monitored orthotopic glioblastoma progression using longitudinal bioluminescence tracking. At Day 9 post-implantation, baseline imaging confirmed identical tumor engraftment and signal intensity across all animals, ensuring uniform baseline metrics before starting treatment (Figure 1A, 1B). By Day 21, the vehicle-treated controls showed a massive, uninhibited surge in bioluminescent signaling, typical of rapid glioblastoma proliferation (Figure 1A). In striking contrast, mice receiving intranasal CBD demonstrated an apparent arrest in signal velocity, successfully containing the tumor footprint (Figure 1A). Quantitative analysis of the photon flux at Day 21 confirmed that this intranasal approach achieved a highly significant suppression of active tumor luminescence compared to vehicle controls (∗∗∗ *P* <0.001; Figure 1B). To bridge these functional imaging patterns with high-resolution structural endpoints, we examined coronal hematoxylin and eosin (H&E) whole-brain sections harvested at Day 21 (Figure 1C). The vehicle control slides revealed large, densely packed glioblastoma masses invading and distorting the cerebral hemispheres (Figure 1C). Conversely, brains from the intranasal CBD cohort presented with significantly smaller, confined lesions that spared the surrounding parenchymal architecture (Figure 1C). Using automated digital threshold tracking to cleanly segment the tissue margins and outline the tumor boundaries (Figure 1D), absolute pixel area quantification (× 1000) verified a major, highly significant contraction in cumulative tumor burden driven by the cannabinoid treatment (*P* <0.01; Figure 1E). Together, these parallel imaging and histological datasets demonstrate that the intranasal CBD platform effectively limits the spatial expansion of glioblastoma.

**Fig. 1.**
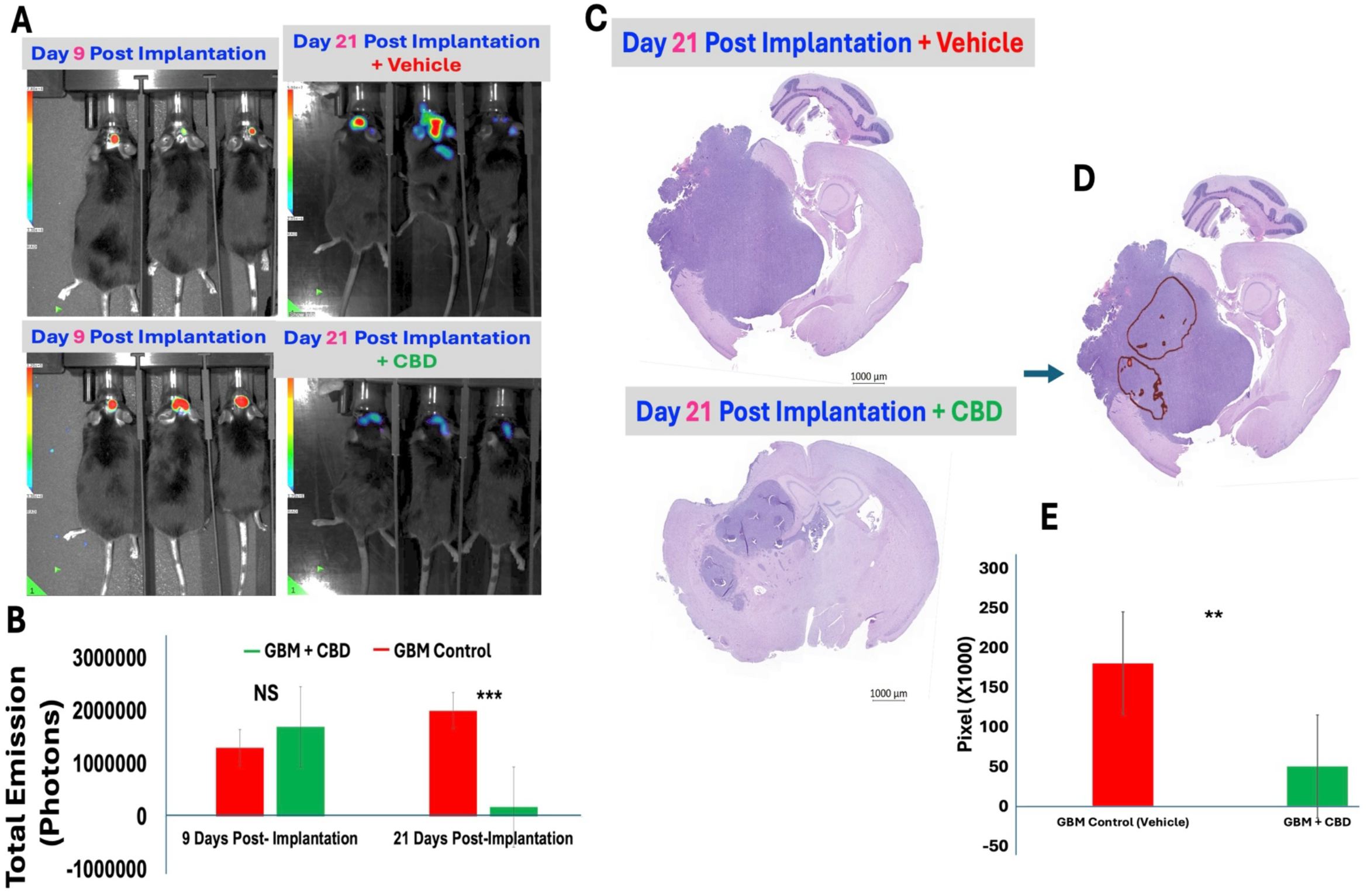
Intranasal delivery of cannabidiol (CBD) limits intraparenchymal expansion of orthotopic glioblastoma (GBM). **(A)** Baseline bioluminescence tracking at Day 9 post-implantation establishes uniform tumor engraftment across cohorts prior to treatment initiation, whereas subsequent imaging at Day 21 reveals uninhibited signal expansion in vehicle controls compared to the marked containment of tumor-derived photon emission in mice receiving intranasal CBD. **(B)** Statistical evaluation of total photon flux confirms these findings, showing a highly significant blunting of tumor signal growth by the terminal checkpoint (∗∗∗ *P* <0.001) despite matching baseline values. **(C)** Whole-slice histological analysis via coronal hematoxylin and eosin (H&E) sections at Day 21 corroborates the in vivo tracking, with vehicle-treated brains exhibiting extensive, space-occupying glioblastoma masses that distort local tissue architecture, while the intranasal CBD cohort shows localized, structurally restricted tumor footprints. **(D)** Digital tissue-boundary threshold tracking systematically maps out the malignant margins to establish objective structural segmentation for area analysis. **(E)** Pixel area measurements (× 1000) quantify these boundaries, verifying a major, statistically significant reduction in absolute neoplastic mass driven by the cannabinoid formulation (*P* <0.01); data are presented as mean ± SD (scale bars: 1000 µm).

### Intranasal CBD Restructures the Intratumoral Microbial Landscape

To investigate how intranasal CBD treatment modifies the tumor-associated microenvironment beyond host immune cells, we characterized the intratumoral microbiome using metagenomic taxonomic profiling. In the Vehicle (Untreated) cohort, analysis revealed a highly diverse and complex baseline microbial community within the glioblastoma parenchyma (Figure 2). This control landscape was defined by a heterogeneous mixture of diverse bacterial lineages, including prominent Actinomycetota, Planctomycetota, and Pseudomonadota networks, along with stable archaeal populations (*Nitrososphaeria*) and a large share of unclassified organisms (Figure 2).

**Fig. 2.**
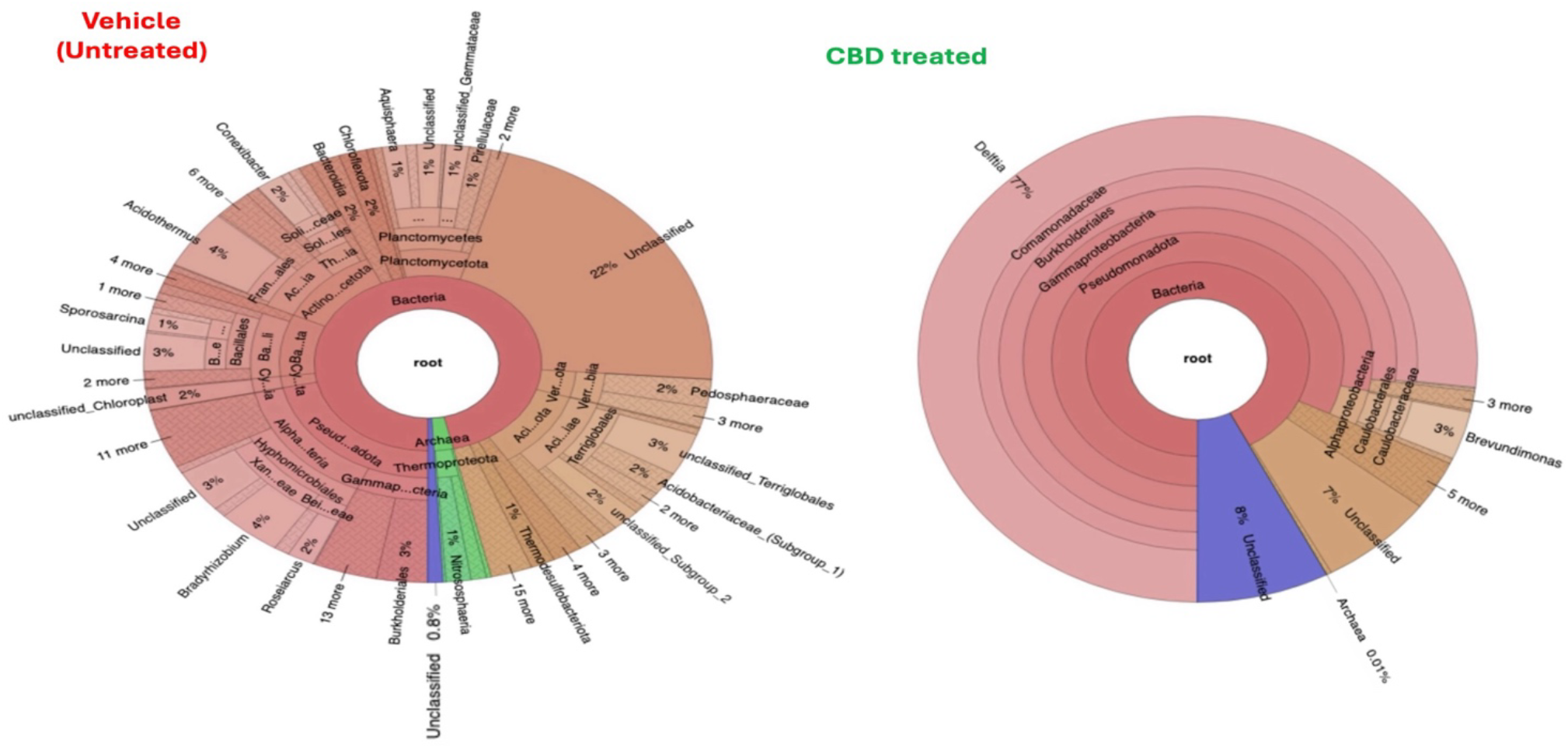
Intranasal cannabidiol (CBD) shifts the intratumoral microbiome profile in orthotopic glioblastoma (GBM). (Vehicle [Untreated]) Taxonomical sunburst profiling of the tumor microenvironment in control mice reveals a highly diverse, heterogeneous microbial ecosystem characterized by a balanced distribution of multiple bacterial phyla, including Actinomycetota, Planctomycetota, and Pseudomonadota, alongside distinct archaeal communities (*Thermoproteota*) and a substantial proportion of unclassified taxonomic groups (22%). **(CBD treated)** Characterization of the microbiome following intranasal CBD therapeutic intervention demonstrates a profound restructuring of the community framework and a dramatic reduction in biodiversity, heavily dominated by a massive clonal expansion of the genus *Delftia* (71%) within the Comamonadaceae family, effectively shifting the tumor-associated microbiome configuration and reducing archaeal abundance to near-detectable limits (0.01%).

Following therapeutic administration, the CBD treated group displayed a striking, systemic rearrangement of this microbial ecosystem (Figure 2). Intranasal CBD effectively collapsed the baseline microbial diversity, shifting the landscape into a highly uniform, low-diversity profile (Figure 2). This treatment-induced selection was dominated by a massive enrichment of the genus *Delftia*, which expanded to comprise 71% of the entire microbial community structure (Figure 2). Concurrently, minor bacterial niches were heavily suppressed, and local archaeal abundance plummeted to a negligible 0.01% (Figure 2). These findings demonstrate that along with suppressing physical tumor progression, the intranasal CBD platform exerts a potent, targeted modulatory effect on the intratumoral microbiome framework.

### Intranasal CBD Bluntly Suppresses Intratumoral Mast Cell and Eosinophil Infiltration

To explore how our formulation influences granular innate immune cell populations within the local tumor landscape, we evaluated the infiltration and distribution profiles of host mast cells and mature eosinophils. Using multi-marker immunofluorescence staining on tumor-bearing brain sections, we observed a massive accumulation of active, mature mast cells (CD117^+^Tryptase^+^) forming prominent clusters at the margins and core zones of the GBM Control group (Figure 3A). This heavy mast cell burden was accompanied by extensive infiltration of double-positive host eosinophils (CD193^+^CD170^+^) throughout the untreated glioblastoma parenchyma (Figure 3B). In striking contrast, the GBM + CBD treatment group displayed a complete disruption of this granulocytic configuration, showing a massive reduction in visible tissue-infiltrating mast cells (Figure 3A) and a near-total loss of the local eosinophil presence (Figure 3B).

**Fig. 3.**
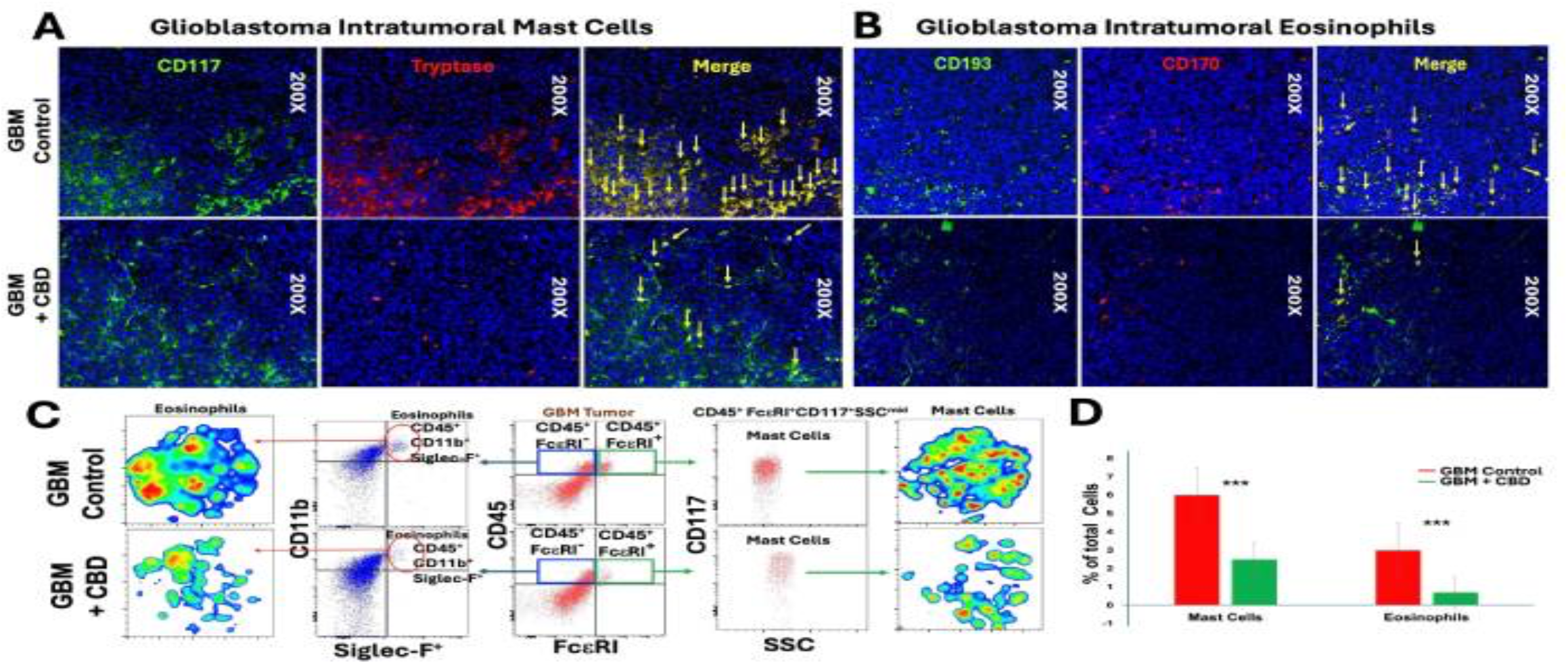
Intranasal cannabidiol (CBD) treatment suppresses the infiltration of host mast cells and eosinophils within the glioblastoma (GBM) tumor microenvironment. **(A)** Representative immunofluorescence panels (200× magnification) of glioblastoma intratumoral mast cells stained for CD117 (green), Tryptase (red), and nuclei (DAPI/blue) show extensive clusters of double-positive mature mast cells (yellow merged signal marked by arrows) in the GBM Control group, which are markedly reduced and scattered following intranasal CBD treatment. **(B)** Representative immunofluorescence panels (200× magnification) of glioblastoma intratumoral eosinophils stained for CD193 (green), CD170 (red), and nuclei (DAPI/blue) demonstrate robust infiltration of double-positive eosinophils (yellow merged signals marked by arrows) in untreated controls, contrasted by a near-complete depletion of these cells in the CBD-treated parenchyma. **(C)** Gating strategies and t-SNE density mapping derived from multicolor flow cytometry analyze the specific granulocyte subsets, tracking eosinophils (CD45^+^CD11b^+^Siglec-F^+^) and mast cells (CD45^+^FcεRI^+^CD117^+^SSC^mid^) to illustrate a comprehensive collapse in both density and total population distribution within the tumor tissue after cannabinoid administration. **(D)** Quantitative comparative evaluation of flow cytometric data confirms that intranasal CBD therapies drive a highly significant, parallel drop in both intratumoral mast cells and infiltrating eosinophils expressed as a percentage of total cells (∗ ∗∗ *P* <0.001); data are presented as mean ± SD.

To securely validate these spatial microanatomical observations with absolute phenotypic quantification, we subjected the processed glioblastoma tissue to multi-parametric flow cytometry and high-dimensional t-SNE density mapping (Figure 3C). Single-cell tracking verified that the dense clusters of tumor-associated eosinophils (CD45^+^CD11b^+^Siglec-F^+^) and regulatory mast cells (CD45^+^FcεRI^+^CD117^+^SSC^mid^) characteristic of the vehicle control baseline collapsed completely into scattered, low-density islands following therapeutic administration (Figure 3C). Statistical analysis of the cumulative single-cell data confirmed that intranasal CBD administration drove a highly significant, parallel drop in both immune cell populations as a percentage of total cells compared to the vehicle group (∗∗∗ *P* <0.001; Figure 3D). These parallel imaging and cytometric datasets demonstrate that the intranasal CBD platform strongly modulates the glioblastoma microenvironment by blocking the influx or survival of tumor-promoting granulocytic lineages.

### Intranasal CBD Induces Ultrastructural Degradation of Tumoral Granulocytes

To explore the microanatomical mechanisms driving the quantitative reduction of tumor-associated granular lineages following therapeutic intervention, we utilized high-resolution transmission electron microscopy (TEM) to evaluate targeted cell-level alterations inside the glioblastoma mass. Under baseline GBM Control (Vehicle) conditions, infiltrating host mast cells exhibited classical healthy morphology, maintaining well-defined cellular boundaries and a cytoplasmic matrix packed tightly with a dense, uniform array of electron-dense secretory granules (Figure 4A). Following therapeutic exposure, profiling of the GBM + CBD cohort revealed profound structural collapse in this population. Intratumoral mast cells captured post-treatment displayed severe degranulation and architectural fragmentation, marked by extensive variations in granule core density, loose structural integrity, and the formation of empty vacuolar networks throughout the lucent cytoplasm (Figure 4B).

**Fig. 4.**
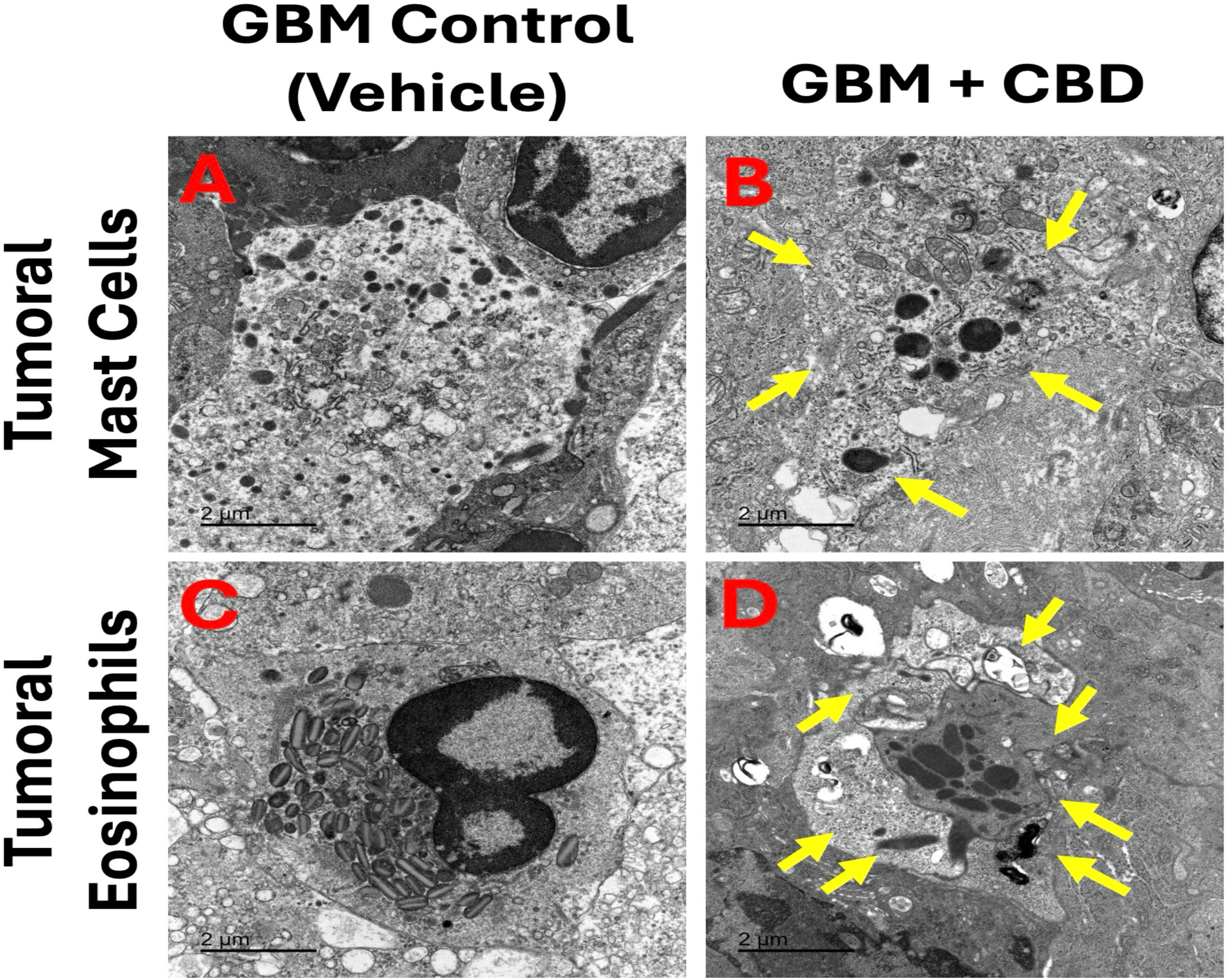
Intranasal CBD delivery alters the ultrastructural integrity of tumor-associated granulocytic lineages. **(A)** Transmission electron micrograph (TEM) of a baseline tumoral mast cell in a vehicle control glioblastoma environment, highlighting a robust, intact cytoplasm heavily laden with a high density of uniform, electron-dense secretory granules. **(B)** TEM of an intratumoral mast cell following intranasal CBD administration, with yellow arrows mapping a distinct therapeutic fingerprint defined by extensive architectural degradation, loose granule remnants with altered core densities, and widespread cytoplasmic clearings. **(C)** TEM of a host eosinophil within the vehicle-treated control parenchyma, exhibiting classic healthy microanatomy characterized by a distinct bilobed nucleus and cytoplasm tightly packed with mature, intact secondary granules containing sharp crystalline cores. **(D)** TEM of an infiltrating eosinophil under intranasal CBD exposure, where yellow arrows document severe structural degeneration, condensing of remaining core granules, necrotic tissue stress, and the prominent development of autophagic clearings filled with dense lipid whorls forming distinct myelin figures. *(Scale bars: 2 µm)*.

Having established the treatment-induced breakdown of the mast cell compartments, we next evaluated the structural response of host eosinophils within the tumor microenvironment. In the vehicle-treated control parenchyma, these infiltrating granulocytes showed no signs of architectural stress, displaying characteristic bilobed nuclear structures and mature secondary granules with intact, signature crystalloid cores (Figure 4C). Conversely, tumor-associated eosinophils subjected to intranasal CBD administration experienced substantial therapeutic stress and degenerative remodeling (Figure 4D). Following treatment, these cells were defined by prominent features of necrotic condensation, severe granule degradation, and a notable development of complex autophagic structures alongside concentric, degenerated lipid membrane whorls forming distinct myelin figures (Figure 4D). Together, these ultrastructural datasets demonstrate that the intranasal CBD platform actively reshapes the local microenvironment by triggering the physical breakdown and structural collapse of tumor-associated granulocytic lineages in a sequential, panel-aligned manner.

## Discussion

The prognosis for glioblastoma (GBM) remains notoriously grim, with standard therapeutic approaches severely limited by restricted drug transport across the blood-brain barrier and an immunosuppressive microenvironment ^21,22^. In this study, we present a paradigm-shifting approach utilizing a non-invasive, locally focused intranasal cannabidiol (CBD) formulation that achieves notable spatial restriction of orthotopic GBM progression. Beyond simple anti-neoplastic activity, our results uncover an intricate dual mechanism of action: a dramatic remodeling of the intratumoral microbiome and a comprehensive, structural deconstruction of tumor-promoting granulocytic lineages. By showing that cannabinoid delivery drives both macro-level tumor containment and micro-level microenvironmental reorganization, these datasets establish a compelling foundation for targeting non-classical immune and microbial axes to manage high-grade malignancies. A central novelty of this work is the comprehensive metagenomic profiling of the intratumoral microbial landscape. While the central nervous system and its tumors were historically regarded as largely immunologically isolated and sterile, accumulating evidence has begun to redefine this traditional paradigm by revealing a far more dynamic microenvironment involving resident immune cells, peripheral immune interactions, and potentially tumor-associated microbial communities ^23-25^. Consistent with this emerging concept, our findings demonstrate that untreated glioblastoma tissue harbors an intrinsic microbial community characterized by distinct Actinomycetota, Planctomycetota, and Pseudomonadota networks, suggesting that microbial ecology may represent a previously underappreciated component of the glioblastoma tumor microenvironment. Crucially, the presence of this diverse, heterogeneous ecosystem, independent of therapeutic intervention, indicates that the intratumoral microbiome plays an active, plausible role in shaping the glioblastoma niche. This complex microbial web may contribute to local tissue remodeling, metabolic signaling pathways, or the maintenance of immune privilege, introducing a highly novel vector in neuro-oncology. Following intranasal CBD therapy, this complex ecosystem underwent a notable rearrangement, collapsing baseline diversity into a highly uniform profile dominated by a 71% clonal enrichment of the genus *Delftia*. This selective shift, accompanied by the near-total loss of local archaeal populations, suggests that our formulation alters the metabolic or structural architecture of the tumor niche, creating a highly selective microenvironment hostile to competitive bacterial strains. Delftia species possess remarkable metabolic versatility, including extensive biotransformation capabilities and the production of bioactive small molecules such as delftibactin and harmane, raising the possibility that they may influence local biochemical signaling within the tumor microenvironment ^26-28^. This treatment-induced selection indicates that intranasal CBD functions as a selective homeostatic modifier. By shifting the microbial equilibrium, the formulation strips the malignancy of its supporting microbial infrastructure, pointing to a novel mechanism where antitumor efficacy is partially mediated through targeted microbiome restructuring.

Beyond microbial modulation, this study introduces a crucial immunotherapeutic paradigm by targeting tumor-associated granulocytes, specifically mast cells and eosinophils. Although the immunobiology of glioblastoma has traditionally been interpreted through the lens of adaptive immunity, tumor-associated macrophages and, more recently, neutrophils, comparatively little attention has been directed toward other granulocyte subsets within the glioblastoma immune ecosystem ^28-31^. Our findings suggest that mast cells and eosinophils may represent previously underrecognized participants in the glioblastoma immune ecosystem, broadening the spectrum of innate immune cells capable of shaping the tumor microenvironment. Our baseline imaging and flow cytometry data reveal a massive influx of active, mature mast cells (CD117^+^Tryptase^+^) and infiltrating host eosinophils (CD193^+^CD170^+^) forming dense clusters throughout the untreated glioblastoma parenchyma. In the untreated tumor, these granulocytes likely act as vital components of the malignancy’s support system, releasing angiogenic factors, matrix metalloproteinases, and immunosuppressive cytokines that facilitate invasion and shield the tumor from surveillance. Intranasal CBD administration drove a complete collapse of this granulocytic framework, transforming these dense cell clusters into sparse, isolated pockets. Our high-resolution transmission electron microscopy (TEM) datasets provide the structural explanation for this depletion, documenting a physical degradation of these populations. Instead of observing normal, healthy cells packed with electron-dense granules, post-treatment granulocytes showed clear signs of therapeutic stress and structural failure. Mast cells exhibited severe degranulation, loss of core integrity, and extensive cytoplasmic clearings. Concurrently, infiltrating eosinophils underwent prominent necrotic condensation and severe granule degradation, marked by the formation of complex autophagic vacuoles and concentric lipid whorls forming distinct myelin figures. This structural deconstruction demonstrates that CBD formulation does not merely prevent granulocyte migration into the central nervous system parenchyma; it actively destabilizes their intracellular machinery, inducing cell-level degradation directly within the tumor niche. This clear target specificity provides a compelling mechanistic explanation for the macro-level tumor restriction observed in our longitudinal in vivo tracking. These findings build upon and extend our established research pipeline demonstrating the multi-faceted therapeutic and protective capacities of CBD within the central nervous system. While our previous work demonstrated that preemptive cannabinoid positioning can temper hyper-inflammatory environments and reinforce local tissue barriers ^17^, we have also successfully demonstrated the efficacy of inhalant CBD in arresting glioblastoma progression by modulating the broader tumor microenvironment ^15^. Furthermore, our exploration of novel biotherapies, including the adoptive transfer of ILC2s, has uncovered distinct tumor-homing capacities that physically interface with the GBM microenvironment ^18^. Consequently, while this is not our first deployment of CBD in an active therapeutic setting against high-grade gliomas, the current study stands as a highly significant translational milestone: it represents the first time a synthetic isolate formulation has been engineered for targeted intranasal administration to selectively trigger structural granulocytic decay. By positioning this specific intranasal cannabinoid formulation at the center of glioblastoma control, this work bridges the gap between conventional oncology, advanced biotherapy, and alternative medicine. Rather than relying on highly toxic chemotherapeutic agents that cause systemic side effects, this approach demonstrates that natural, modified plant-derived compounds can be delivered via non-invasive routes to modify local tissue architecture. This has significant implications for alternative and complementary medicine, providing a rigorous, mechanism-driven scientific foundation for a class of compounds frequently used empirically but rarely validated with such high-resolution ultrastructural and metagenomic evidence. Furthermore, this granulocytic and microbial disruption opens up exciting avenues for modern combination therapy. Because the intranasal CBD platform targets pathways entirely separate from standard DNA-alkylating agents or anti-angiogenic therapies, it represents an ideal partner for multimodal protocols. Combining standard-of-care treatments with an intranasal formulation that dismantles granulocytic protection and resets the intratumoral microbiome could prevent the emergence of therapeutic resistance, offering a highly comprehensive approach to treating high-grade gliomas. From a clinical and translational perspective, introducing granulocytic and microbial modulation as a therapeutic target addresses several long-standing bottlenecks in neuro-oncology. First, the intranasal route represents a patient-friendly delivery system that bypasses systemic hepatic metabolism, maximizes local tissue availability within the central nervous system, and avoids the complications associated with invasive intracranial delivery methods. This simple, non-invasive translation can drastically improve drug availability and patient compliance, allowing for consistent long-term management outside a hospital setting. Second, by identifying tumor-associated mast cells and eosinophils as crucial players in glioblastoma progression, this study provides a novel therapeutic target that can be readily evaluated using existing clinical protocols. Strategies aimed at monitoring and systematically disrupting these granulocytic populations can be integrated into current treatment regimens to improve overall survival rates and minimize cognitive decline. Ultimately, by combining non-invasive delivery with multi-target microenvironmental disruption, this platform offers a viable path toward transforming glioblastoma from a rapidly terminal diagnosis into a manageable condition, significantly improving long-term outcomes and quality of life for oncology patients.

In conclusion, this study shatters the traditional, narrow focus of neuro-oncology by unveiling an entirely unchartered therapeutic axis in the fight against glioblastoma. By engineering a novel, non-invasive intranasal synthetic CBD isolate, we have demonstrated for the first time that aggressive intracranial progression can be arrested not merely by attacking the tumor cells themselves, but by systematically dismantling the structural and microbial ecosystems that sustain them. Our discovery that glioblastoma progression relies on a complex, baseline intratumoral microbiome, and that this landscape can be targeted and collapsed, redefines our understanding of the glioma niche. Coupled with the striking, high-resolution visual evidence of physical degradation inside tumor-associated mast cells and eosinophils, these findings elevate granulocytic and microbial modulation from an academic concept to an active therapeutic reality. By bridging advanced biotherapy with an elegant, patient-ready delivery platform, this work opens a new horizon for non-toxic, multi-modal oncology, offering a sophisticated and highly effective strategy to break therapeutic resistance, optimize drug availability, and fundamentally rewrite the clinical outcome for patients facing a high-grade malignancy.

## Acknowledgements

Authors are thankful to Medicinal Cannabis of Georgia for providing help in optimizing the CBD dosage.

## Declaration of AI Assistance

AI-based software (e.g., ChatGPT) was used solely for language refinement and grammar editing; all scientific content and interpretation were entirely developed by the authors.

## Data Availability Statement

The original contributions presented in this study are included in the article. Further inquiries can be directed to the corresponding authors.

## Author Contributions

Lei P Wang (Conceptualization, Data curation, Formal analysis, Funding acquisition, Investigation, Methodology, Resources, Supervision, Visualization, Writing—original draft, Writing—review & editing); Bidhan Bhandari (Data curation, Visualization, Investigation); Sahar Emami Naeini (Data curation, Visualization, Investigation); James T Earwood (Data curation, Investigation); Brendan Marshall (Data curation, Visualization); Chandramohan Wakade (Writing—review & editing), Jack C Yu (Methodology, Investigation, Writing—review & editing); Ali S Arbab (Resources, Writing— review & editing); Évila Lopes Salles (Investigation, Formal analysis); and Babak Baban (Conceptualization, Data curation, Formal analysis, Funding acquisition, Investigation, Methodology, Project administration, Resources, Supervision, Visualization, Writing—original draft, Writing—review & editing).

